# Identifying tumor cells at the single cell level

**DOI:** 10.1101/2021.10.15.463909

**Authors:** Jan Dohmen, Artem Baranovskii, Jonathan Ronen, Bora Uyar, Vedran Franke, Altuna Akalin

## Abstract

Tumors are highly complex tissues composed of cancerous cells, surrounded by a heterogeneous cellular microenvironment. Tumor response to treatments is governed by an interaction of cancer cell intrinsic factors with external influences of the tumor microenvironment. Disentangling the heterogeneity within a tumor is a crucial step in developing and utilization of effective cancer therapies. The single cell sequencing technology enables an effective molecular characterization of single cells within the tumor. This technology can help deconvolute heterogeneous tumor samples and thus revolutionize personalized medicine. However, a governing challenge in cancer single cell analysis is cell annotation, the assignment of a particular cell type or a cell state to each sequenced cell. One of the critical cell type annotation challenges is identification of tumor cells within single cell or spatial sequencing experiments.This is a critical limiting step for a multitude of research, clinical, and commercial applications. A reliable method addressing that challenge is a prerequisite for automatic annotation of histopathological data, profiled using multichannel immunofluorescence or spatial sequencing. Here, we propose Ikarus, a machine learning pipeline aimed at distinguishing tumor cells from normal cells at the single cell level. We have tested ikarus on multiple single cell datasets to ascertain that it achieves high sensitivity and specificity in multiple experimental contexts.

## Introduction

Cancer is a disease that stems from the disruption of cellular state. Through genetic perturbations, tumor cells attain cellular states that give them proliferative advantage over the surrounding normal tissue [1]. The inherent variability of this process has hampered efforts to find highly effective common therapies, thereby ushering the need for precision medicine [2]. The scale of single cell experiments is poised to revolutionize personalized medicine by effective characterization of the complete heterogeneity within a tumor for each individual patient [3,4].

Recent expansion of single cell sequencing technologies have exponentially increased the scale of knowledge attainable through a single biological experiment [5]. The information contained within a single high throughput single cell experiment enables not only characterization of variable stable states (i.e. cell types, and cell states), but also functional annotation of individual cells, such as prediction of the differentiation potential, susceptibility to perturbations, and inference of cell - cell interactions [6].

As with all new technologies, high throughput single cell sequencing also created new computational challenges [7]. One outstanding problem in single cell data analysis is cell annotation - assignment of a particular cell type or a cell state to each sequenced cell. The size of the generated datasets made manual annotation approaches utterly unfeasible, while the peculiarities of data generation prompted the development of novel innovative classification methods [8–13].

Currently, there are three general types of cell annotation tools: manifold matching software, which aligns the shape of an unlabeled dataset to an expertly labeled dataset [14], deep learning based tools which try to model the batch effects through latent space embedding [15–17]; and gene set based classifiers that use known marker genes for cell type assignment [8]. Although a multitude of methods have recently been developed for single cell data annotation, most of them have not been evaluated for robustness. Most often, it is unclear whether the data used for training, validation, and testing are independent. It is not evident whether the learned model will perform equally well on the data profiled using different sequencing technologies (i.e. Drop-Seq, 10x, CEL-seq or Fluidigm C1), produced by different laboratories, or originating from a different biological source (i.e. same cell types, coming from different mice or humans). The sensitivity of the single cell sequencing methods comes with a price - the generated data contain the information about the interesting biology mixed with unwanted variability [18]. The sources of this variability have still not been comprehensively characterized, and can not be modeled explicitly [19]. Therefore, special care needs to be taken when developing machine learning methods on single cell datasets, that the learned association really is between biological variables, and is not confounded by the properties of the data.

We set out to answer a simple question: “Is it possible to make a classifier that would correctly differentiate tumor cells from normal cells?”. The utility of such a tool would be manyfold. Automatic characterization of tumor cells can be used for automation of neoantigen prediction [20,21]. It would enable automatic characterization of tumor cell states, such as tumor stem cells [22]. Computational delineation of cancer cells would greatly expedite biomarker discovery, and the development of companion diagnostic tools for measuring drug efficacy, during preclinical and clinical drug development. Last but not least, such a tool can be used for automatic annotation of histopathological data, profiled using multichannel immunofluorescence or spatial sequencing [23].

We have built Ikarus, a stepwise machine learning pipeline for tumor cell classification. Ikarus consists of two steps: 1) discovery of a comprehensive tumor cell signature in the form of a gene set by consolidation of multiple expertly annotated single cell datasets. 2) training of a robust logistic regression classifier for stringent discrimination of tumor and normal cells followed by a network-based propagation of cell labels using a custom built cell-cell network [24]. With the goal of developing a robust, sensitive, and reproducible *in silico* tumor cell sorter, we have tested Ikarus on multiple single cell datasets of various cancer types, obtained using different sequencing technologies, to ascertain that it achieves high sensitivity and specificity in multiple experimental contexts. We have strictly adhered to machine learning best practices, to avoid results contamination by information leaking from training into testing.

## Results

### Identification of a robust marker gene set

Cell type annotations in any particular experiment are inherently noisy. This is partly due to the properties of single cell data, such as the different number of detected genes in each cell, the influence of sample processing, and our limited knowledge of biomarkers that are necessary for comprehensive annotation of cell types and cell states. We hypothesized that we can find robust markers of cellular states by comparing multiple independent annotated datasets from diverse tumor types.

We have integrated datasets from colorectal [25] and lung [26] cancers that were profiled using 10x single cell droplet technology. The authors of both studies have annotated each cell in the dataset with the corresponding cell types - including whether the cells are healthy or cancerous. We have employed a two step procedure to find tumor-specific gene markers. First, using differential expression analysis, we selected genes that are either enriched or depleted in cancer cells per dataset (See Methods). To obtain the final gene list, we took an intersection of the gene sets from each of the datasets (Figure 1A). We ended up with a set of 162 genes that were significantly enriched in cancer cells across multiple datasets (Table S1 - Sheet 1). The resulting set of genes showed high specificity for cancer cells, from the head and neck cancer samples [27] (Figure 1C). This result indicates that the gene set contains information relevant for discriminating tumor cells from non-tumor cells in multiple different tumor types.

**Figure 1.**
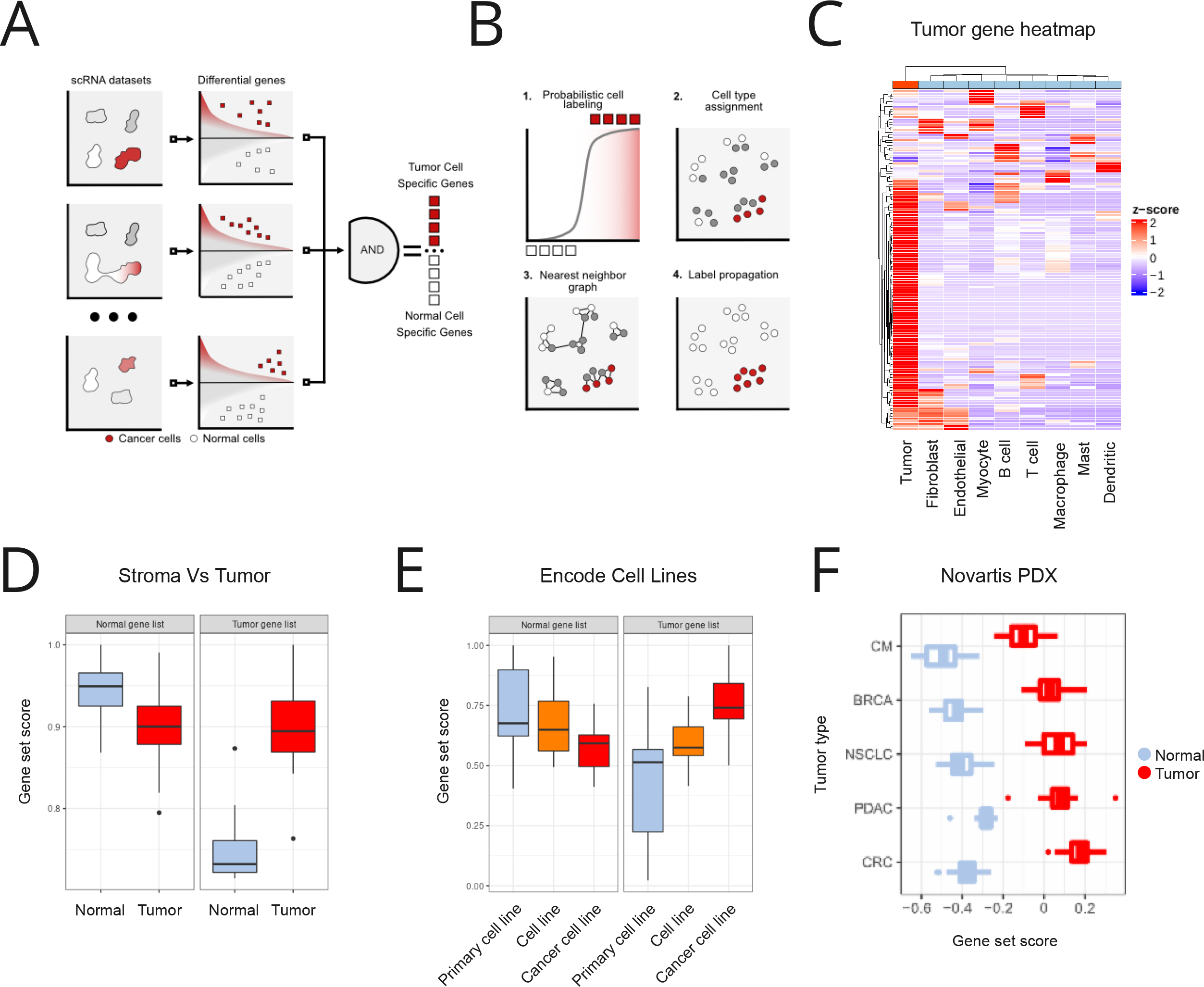
Integration of multiple datasets enables robust extraction of informative gene sets. A-B) Ikarus workflow. Ikarus is a two step procedure for classifying cells. In the first, integration of multiple expert labeled datasets enables the extraction of robust gene markers. The gene markers are then used in a composite classifier consisting of logistic regression and network propagation. C) The heat map depicts the expression of the tumor specific genes in the Tirosh head and neck cancer dataset [27], which was not used for the gene list definition. Out of the 162 genes from the list, 132 were found to be expressed in the Tirosh dataset. The gene list contains two sets of genes: genes that are highly enriched in tumor cells compared to all other cells and genes that are highly enriched in tumor cells compared to each individual cell type. D) Comparison of gene signature scores in laser microdissected gastric cancer data [28]. The normal gene list shows significantly lower signature scores in cancer samples, when compared to the cancer-associated normal tissue. The tumor gene signature is significantly higher for cancer samples than the normal tissue. E) Primary cells and cancer cell lines have significantly different gene signature distributions. The normal-cell gene signature shows a gradual reduction in gene signature score distribution when compared in primary cells, cell lines, and tumor cell lines. The gene signature shows the complete opposite effect. Cancer cell lines have the higher gene signature score distribution, followed by cell lines, and primary cells. F) Patient derived xenografts (PDX) show significantly higher tumor gene signature score, than the normal gene signature score. The same pattern is observed in multiple cancer types.

The same procedure was applied to the healthy cell types. We extracted genes enriched in each cell type, when compared to the tumor cells. The resulting gene set was then merged between multiple datasets. This “normal” cell gene signature contains both cell type specific markers, and genes which are specifically depleted in the tumor cells (Figure S1A). To validate the specificity of the novel tumor specific gene set, we have analysed a gastric cancer dataset [28], where multiple areas of cancer and cancer-associated normal cells were separated using laser-capture microdissection (LCM) and profiled using RNA sequencing. Using normal and tumor gene signatures that were identified by Ikarus, we have scored the tumor and the associated normal cells. As expected, dissected sections coming from the cancerous lesions had significantly higher median tumor score than the surrounding normal tissue (Figure 1D, right panel). In line with the latter, normal tissue scored higher than cancerous lesions when the normal gene signature was scored (Figure 1D, left panel).

As another line of evidence, we have downloaded the expression data for primary, normal, and cancer cell lines from the ENCODE database [29,30] (See Methods). Tumor signature scores were on average highest in cancer cell lines, diminished in normal stable cell lines, reaching its lowest average in primary cells (Figure 1E, left panel). When scoring using the normal (non-cancer) cell signature, an opposite trend was observed, i.e. score was highest in primary cells, intermediate in normal stable cells and the lowest in cancer cell lines (Figure 1E, right panel).

Further, we tested the discriminatory power of the normal and tumor gene lists in multiple cancer types. To this end we have used the patient-derived xenograft (PDX) samples from five cancer types provided by [31], and all of the cancer cell lines provided by the cancer cell line encyclopedia (CCLE) [32]. The tumor signature score was significantly higher than the normal signature score in all PDX cancer types (Figure 1F) and all cancer cell lines screened in CCLE (Figure S1B). Surprisingly, the tumor signature list produced significantly reduced scores for cell lines stemming from blood related cancers (LAML, CLL, LCML, MM, DLBC).

### Accurate delineation of cancer cells

In the first step of classification, Ikarus derives the tumor and normal gene set scores. The tumor and normal gene set scores are then used in a logistic regression classifier, to delineate cells with high probability of being tumorous or normal. The classification step is followed by the propagation of the cancer/normal label through a custom based cell - cell network (Figure 1B). The cell - cell network is derived from the same gene sets that are used for robust scoring. By using only tumor or cell type specific genes, the resulting network separates communities that represent either tumor or normal cell states.

We trained Ikarus on scRNA-seq datasets of colorectal [25] and lung carcinomas [26]. The algorithm was further tested on the following cancer types: lung [33], head & neck cancer [27], and neuroblastoma [34]. Figure 2A shows the performance of Ikarus classification on all of the test datasets. Ikarus achieves an average balanced accuracy of 0.98 which is substantially higher than other classical machine learning methods. In addition to the standard machine learning methods (SVM, random forest, and logistic regression), we have compared Ikarus to the top ranking tailored cell type classifiers, as evaluated in [8]: SingleCellNet [9] and ACTINN [35]. Ikarus again consistently showed better performance discriminating tumor cells from normal cells.

**Figure 2.**
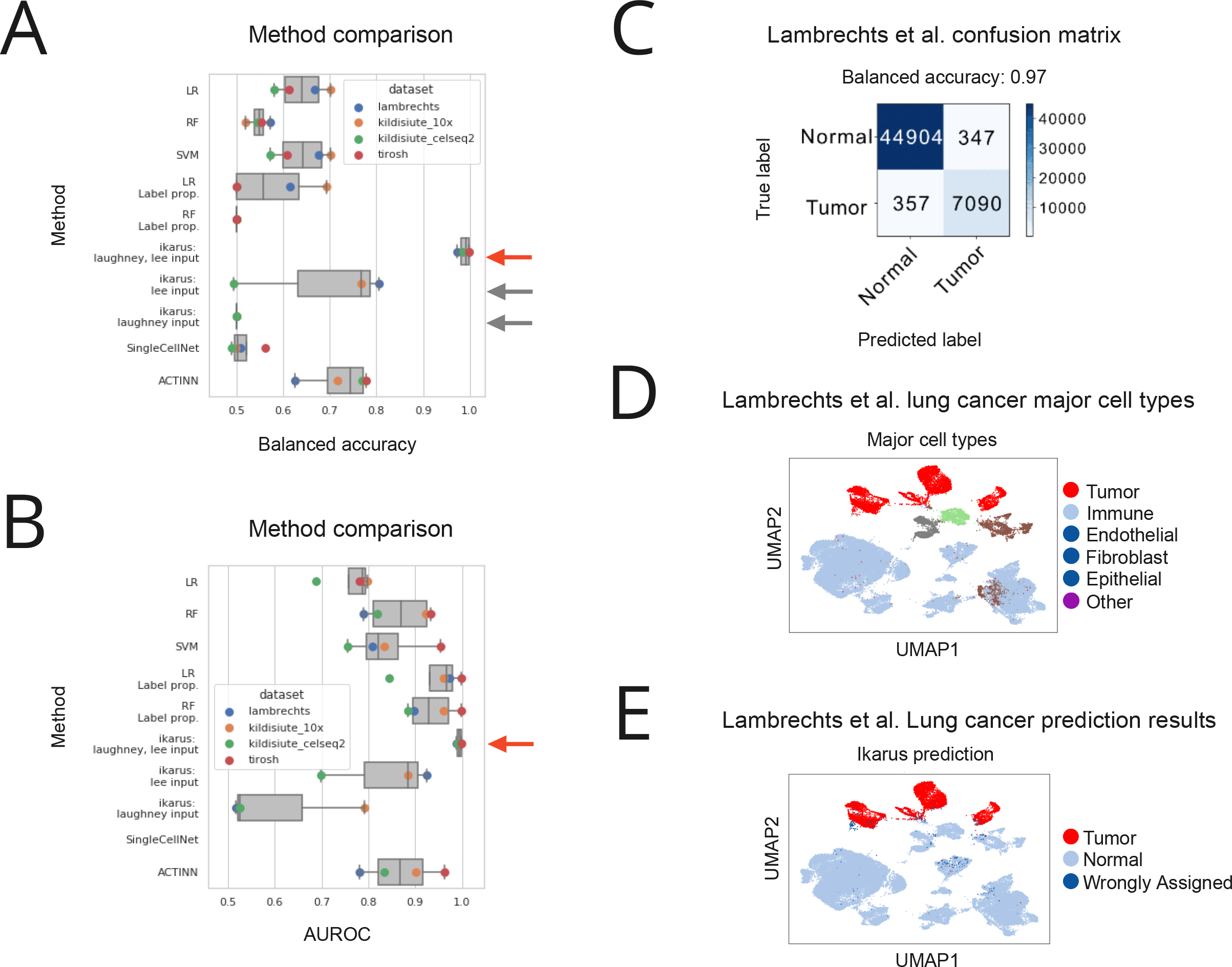
Ikarus accurately classifies tumor and normal cells. A) Balanced accuracy for classification of tumor and normal cells, for each of the test data sets. Red arrow highlights performance of Ikarus classifier. The grey arrows highlight performance of Ikarus, but trained on only a single data set (either Lee et al. or Laughney et al.). B) Area under receiver operating characteristic (AUROC) performance for each classifier. C) Confusion matrix showing the performance of Ikarus classifier on the Lambrechts lung cancer dataset. D) Cell type annotation of the Lambrechts et al. dataset. E) Lambrechts dataset labeled by Ikarus classifier.

We have chosen balanced accuracy as a measure of performance because of the large imbalance of classes. The datasets contained, on average, 7 times more normal cells than annotated cancer cells (Table S1 - Sheet 2). To give an unbiased view on the performance, Figure 2B shows the area under the receiver operating characteristic (AUROC) distribution for the different datasets. Ikarus also achieves a higher average AUROC than other methods. Having a high AUROC value, and low balanced accuracy is an indication of class imbalance. The classification error of the classical machine learning methods, having high AUROC and low balanced accuracy, is not uniformly distributed - they struggle with a high false positive rate. We were wondering whether Ikarus could be trained using just one dataset. When Ikarus was trained separately on colorectal [25], or the lung cancer [26] datasets, the performance dropped substantially (Figure 2A, grey arrows). Figure 2C shows the classification accuracy for the Lambrechts lung cancer dataset [33]. The Lambrechts dataset was not used for training nor gene signature definition. Figure 2D and E show the classification accuracy overlaid on UMAP [36] embeddings of the Lambrechts lung cancer dataset [33]. Ikarus correctly classifies normal cells, irrespective of the underlying cell types, while the misclassified cells are uniformly distributed between all of the cell types (Table S1 - Sheet 5). The erroneous classifications are equally distributed between false positives and false negatives (UMAPs for other datasets are reported in Figure S2).

In order to test the robustness of Ikarus across different single cell sequencing technologies, we applied Ikarus on a dataset of neuroblastoma tumors sequenced by both 10X genomics and CEL-Seq2 protocols by Kildisiute et al. [34]. Ikarus achieved a high classification accuracy (balanced accuracy of 0.98) on both datasets, irrespective of the profiling technology (Figures S2B and S2C). The false positive rate we observed in the test datasets (1 - 3%) can be partly attributable to occasional erroneous labeling of cells by the authors of the corresponding studies. The lack of a perfectly labeled single cell tumor sequencing dataset makes it difficult to quantify the actual rate of false positive predictions by our method. One possible strategy to remedy this issue is to test our method on a dataset that is presumably free of tumor cells, in other words, a healthy tissue sample. To ascertain the actual false positive rate for tumor cell classification, we have tested Ikarus on the single cell data from peripheral blood of a healthy individual [37], where all cells are expected to be non-tumorous. Ikarus labeled all cells as non-tumorous (Figure S2D).

Further, we were interested in how the accuracy of the classification changes in regards to the size and structure of the gene set. First, we have conducted an ablation study, where we removed from 20% to 80% of randomly selected genes from the list (Figure 3A). The removal of up to 40% of the genes from the list leads to a ∼12% (from 99% to 87%) drop in median accuracy. If 80% of the gene list is removed, the classification becomes random (median accuracy tends to ∼50%).

**Figure 3.**
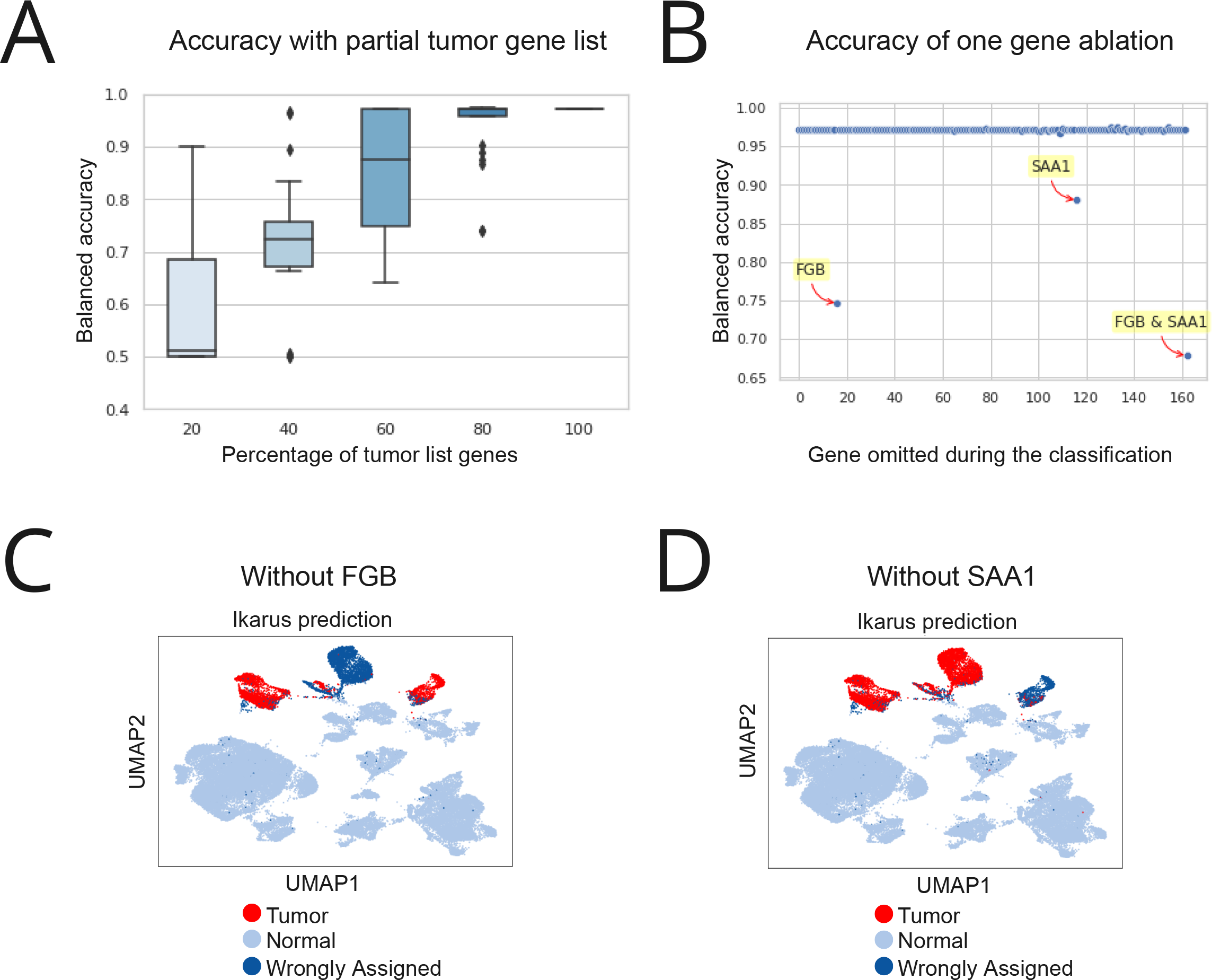
Ikarus performance under perturbation. A) Performance of Ikarus classifier when 20% to 80% of the input list is removed. B) Performance of Ikarus classifier upon single gene removal from the gene list. 160 out of the 162 genes were inconsequential for the classification. FGB (fibrinogen) and SAA1 (serum amyloid alpha) ablation significantly reduced the classification accuracy, but only on the Lambrechts et al. lung cancer dataset. C-D) FGB and SAA1 are respectively strong markers for one of the cell state clusters in the lung cancer data set.

Next, we explored how much the accuracy of the classification depends on individual genes. To test this, we sequentially removed each individual gene from the set and repeated the classification. For 160 out of the 162 genes, there was no observable change in the classification accuracy on the test data sets (Figure 3B). The accuracy on the Lambrechts lung cancer [33] dataset was, however, particularly sensitive to the omittance of two genes: serum amyloid A (SAA1), and fibrinogen beta chain (FGB) (Figure 3C). Each gene is a marker for a tumor specific cell cluster in the Lambrechts dataset (Figures 3C and 3D), and their removal influences the classification probability of cells constituting that particular cluster. Such dependence was not observed for other test data sets (Figure S3A).

### Properties of the tumor gene signature

Having observed the high accuracy performance of Ikarus based on the detected tumor gene signature, we ventured forth to obtain a deeper characterisation of the functional content of these genes. Specifically, their involvement in the development of cancer and their roles in the prognosis for the patients.

Firstly, we were interested whether the genes within the tumor gene signature conform into expression modules, or whether their expression distribution is independent. We calculated the pairwise Pearson correlation between the genes from the signature for all datasets. To our surprise, the correlation between the genes was largely zero (Figure 4A). We found only a single module (containing 34 genes) that was robustly present in all datasets (Figure S4A). Genes in this module are annotated as belonging to the cell cycle. We further inspected whether the classification accuracy depends on these cell-cycle related genes. The removal of the 34 cell cycle-related genes did not affect the classification accuracy (results not shown). The tumor gene signature had, to our surprise, little overlap with established cancer related gene sets. When compared with the gene sets annotated in the CancerSEA database of cancer functional states [38], our tumor gene list had zero or very few overlaps with most CancerSEA gene sets, except for the cell cycle genes, which shared only 9 genes with our tumor gene list (Figure 4B). Co-expression analysis, using SEEK [39], again showed that the tumor gene signature is partially related to cell cycle, and DNA replication (Figure 4C). In addition, we saw no overlap with the cancer hallmarks from MSIGDB [40] (Figure S4B). When compared to the complete MSIGDB database, the tumor gene signature preferentially overlapped with the cell cycle hallmark (Figure 4D). Gene ontology (GO) analysis using gprofiler2 [41] showed an enrichment of terms exclusively related to cell cycle and mitosis (Figure S4C).

**Figure 4.**
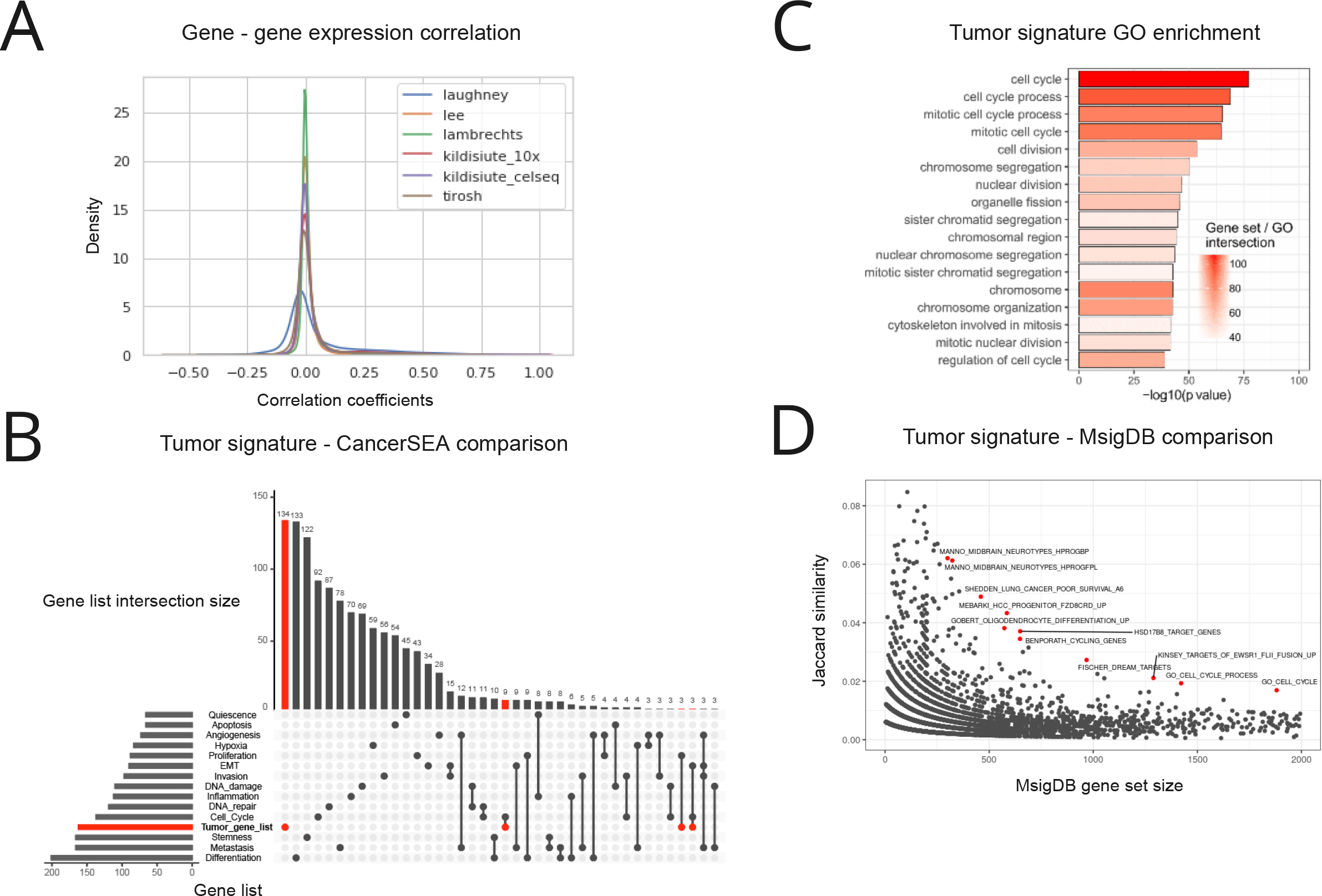
Properties of the tumor gene signature. A) Tumor gene signature co-expression analysis in tumor cells. Co-expression is measured as Pearson correlation between all pairs of genes from the gene list. B) Tumor gene signature shows little overlap with known cancer-associated gene sets. The tumor gene signature was compared to the CancerSEA database. Out of the 162 genes, 134 showed no overlap with any of the gene lists. 9 genes overlapped with the cell cycle gene list. Only intersections of size 3 and more are shown. C) Results of the gene coexpression analysis using SEEK. D) Tumor gene signature shows limited overlap with most of the gene sets from MsigDB. The most enriched gene set corresponded to the cell cycle.

The enrichment of cell cycle and DNA replication related functional terms in our tumor gene set (Figure S4C) led us to hypothesize that the novel gene set differentiates promptly cycling cells. To test this hypothesis, we inspected the correlation of the tumor gene set scores with the growth rates detected in Patient Derived Xenograft (PDX) samples from [31] and the doubling times of the cancer cell lines from CCLE [32]. Unexpectedly, there was no correlation between the tumor signature score and the PDX growth rates in any of the reported cancer types (Figure S4D). Repetition of the analysis on the cell line doubling times from CCLE again revealed the same lack of correlation (Figure S4D).

We were interested in whether the tumor cell signature is predictive of survival outcomes in cancer. From the protein atlas database (http://www.proteinatlas.org) [42] we extracted genes predictive of survival in one or more cancers. The overlap of tumor gene signature with the extracted gene set showed that more than 75% of tumor signature genes are predictive of unfavourable prognosis in at least one cancer type (Figure 5A). Interestingly, when stratified by cancer type, tumor signature genes reported to be unfavorable are overrepresented among 5 cancer types: liver, renal, pancreatic, lung and endometrial cancers (Figure 5B). An analogous analysis was done with data taken from [43], where the authors systematically calculated the risk predictive status for all genes in TCGA cancer types. The analysis showed that the cancer specific genes have a significantly higher Stoufer’s Z value (a measure of how significantly the gene expression predicts the risk status in any cancer) than the rest of the annotated gene set (Figure 5C). Furthermore, the same trend was observed in 22 out of 33 profiled cancer types (Welch two sample t-test, Bonferroni adjusted p-value < 0.05) (Figure 5D).

**Figure 5.**
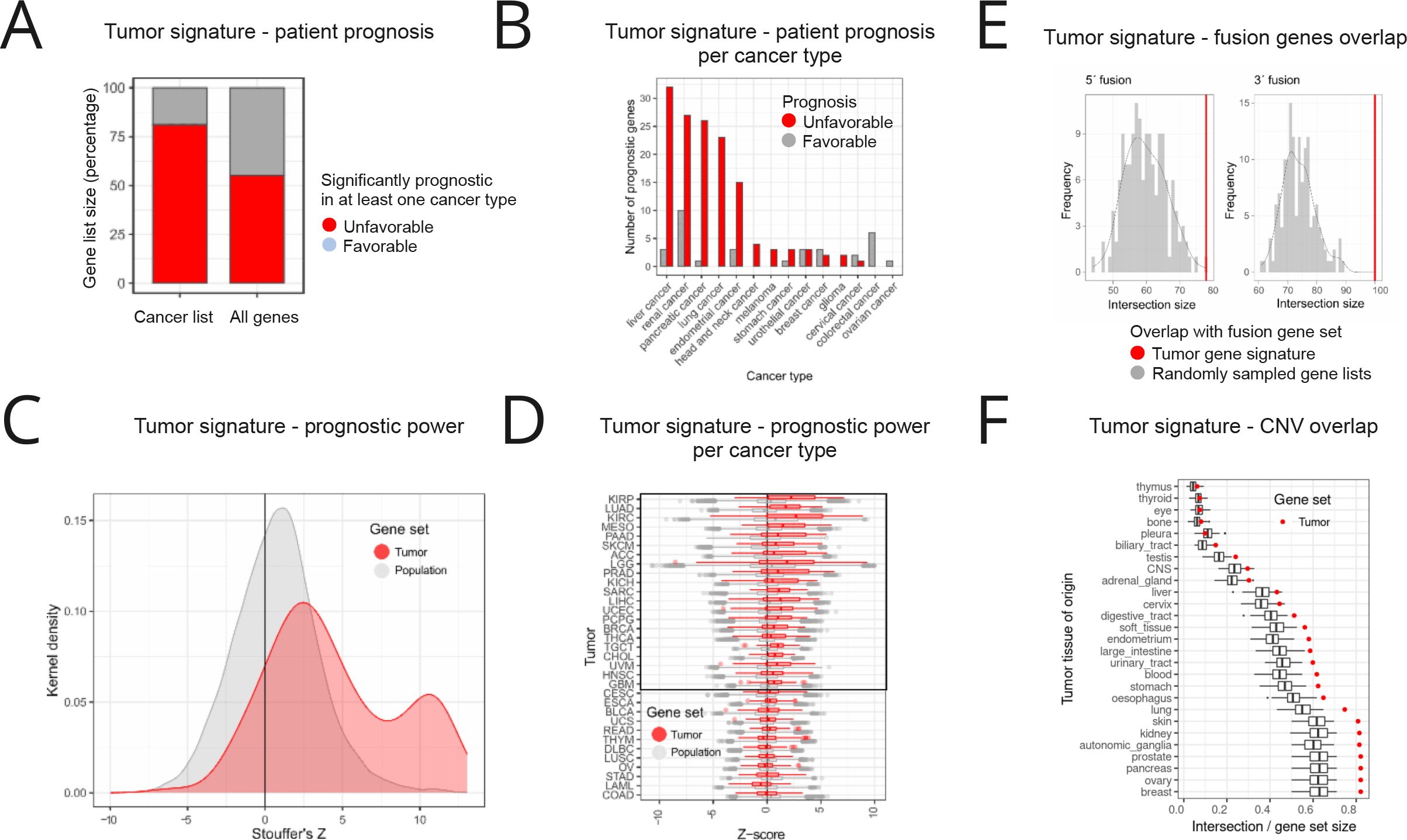
Tumor gene signature is predictive of survival and associated with copy number variations. A) Tumor gene signature genes are more likely to be significantly prognostic for patient survival in at least one cancer type. Data was extracted from the protein atlas. B) The association of tumor gene signature genes with poor survival outcomes is evident in multiple cancer types. C) Data from [43], show that the tumor gene signature genes have much higher Stoufer’s z (association with poor survival outcomes) than rest of protein coding genes. D) Increased association of the tumor signature genes with negative survival is apparent in 22 out of 33 cancer types. E) Genes from the tumor gene signature are more likely to participate in 3’ or 5’ fusions. When compared to sets of randomly drawn genes, the tumor signature genes had a significantly higher probability of participating in genomic fusions. The red vertical line depicts the overlap of the tumor signature list with the corresponding gene fusion list. F) Genes from the tumor gene signature are frequently found in amplified or deleted genomic regions. We have measured the percentage of the gene list which overlaps with the known CNV regions for each cell line in the CCLE dataset. Background distributions were derived from expression matched randomly sampled gene lists.

Next, we wanted to explore how often the genes from the tumor gene list participate in genomic rearrangements, particularly gene fusions, which are frequent drivers of oncogenic events in multiple cancer types. We downloaded the known cancer gene fusions from the ChiTaRS [44] database, and inspected the overlap with the novel cancer defining gene list. To establish enrichment, we compared the overlap with a background consisting of random gene sets. Genes from the tumor gene signature have a significantly higher probability of participating in both 3’ and 5’ fusions, than a random set of genes (Figure 5E).

Gain and loss of DNA content is a ubiquitous property of tumor cells. Copy number variation (CNV) profiles that arise from genomic rearrangements create unique genomic signatures that can be used for characterization, and discrimination of different tumor types [45]. We wondered how prevalent genes from the tumor gene signature are in the known CNV regions. To this end, we have compared the intersection of the tumor signature list with the CNV data from CCLE. We compared the tumor gene list intersection to a background distribution constructed by randomly sampling expression matched gene sets. The tumor gene list had a significantly higher overlap with the known CNV regions in the majority of profiled cancer types (Figure 5F) irrespective of CNV frequency.

### Multi-omics analysis reduces the false positive rate of classification

Characterization of biological systems from multiple viewpoints often produces synergistic insights into the underlying biology. We wondered whether the classification accuracy of Ikarus algorithm can be improved by using multi-omics measurements. To this end, we have used the inferCNV [46] to extract copy number variations (CNVs) from the single cell RNA sequencing data. InferCNV is a Bayesian method, which agglomerates the expression signal of genomically adjointed genes to ascertain whether there is a gain or loss of a certain larger genomic segment. We have used inferCNV to call copy number variations in all samples used in the manuscript.

Firstly, we wondered whether the copy number variations could be used as universal markers for discriminating between normal and cancer cells. We trained random forest classifiers to discriminate between the expert labeled normal and tumor cells. One classifier was trained on each sample. Each of the classifiers was tested on all samples. As expected, when evaluated on the sample which was used for the training, each random forest classifier correctly discriminated between the cancer and tumor cells (Figure 6A). The classifiers, however, did not generalize to other cancer types - they all suffered from a high false positive rate. We tried to improve the generalization of the classification by training on multiple datasets. We trained a random forest classifier on joint Lee et al. and Laughney et al. data, and tested on all other datasets. Using multiple datasets for training did not improve the results of the classification on out of sample cells (Figure 6B). We then wondered whether the CNV calls could be used in conjunction with the gene expression data to improve Ikarus classification of tumor and normal cells. We looked at the average CNV value and the variance of CNV values in cells, which were misclassified by Ikarus in data from Lee et al. and Laughney et. al. Both the average CNV value and the variance of CNVs were significantly higher in cancer cells, which were misclassified as normal cells (Figure 6C). This indicated that by integrating the CNV scores with the gene expression classifier, we might increase the classification accuracy. We have added an additional proofreading step into the classification procedure. We trained a logistic classifier on inferred CNVs, with Ikarus predicted cell type labels as the dependent variable. Cells which obtained highly probable discordant class labels from the CNV classifier, had their labels flipped. Using the proofreading step, the average balanced accuracy stayed the same for all of the samples. We have however noticed a sudden drop in the false positive rate, with a marginal increase in the false negative rate (Figure 6D, Table S1, Sheet 3).

**Figure 6.**
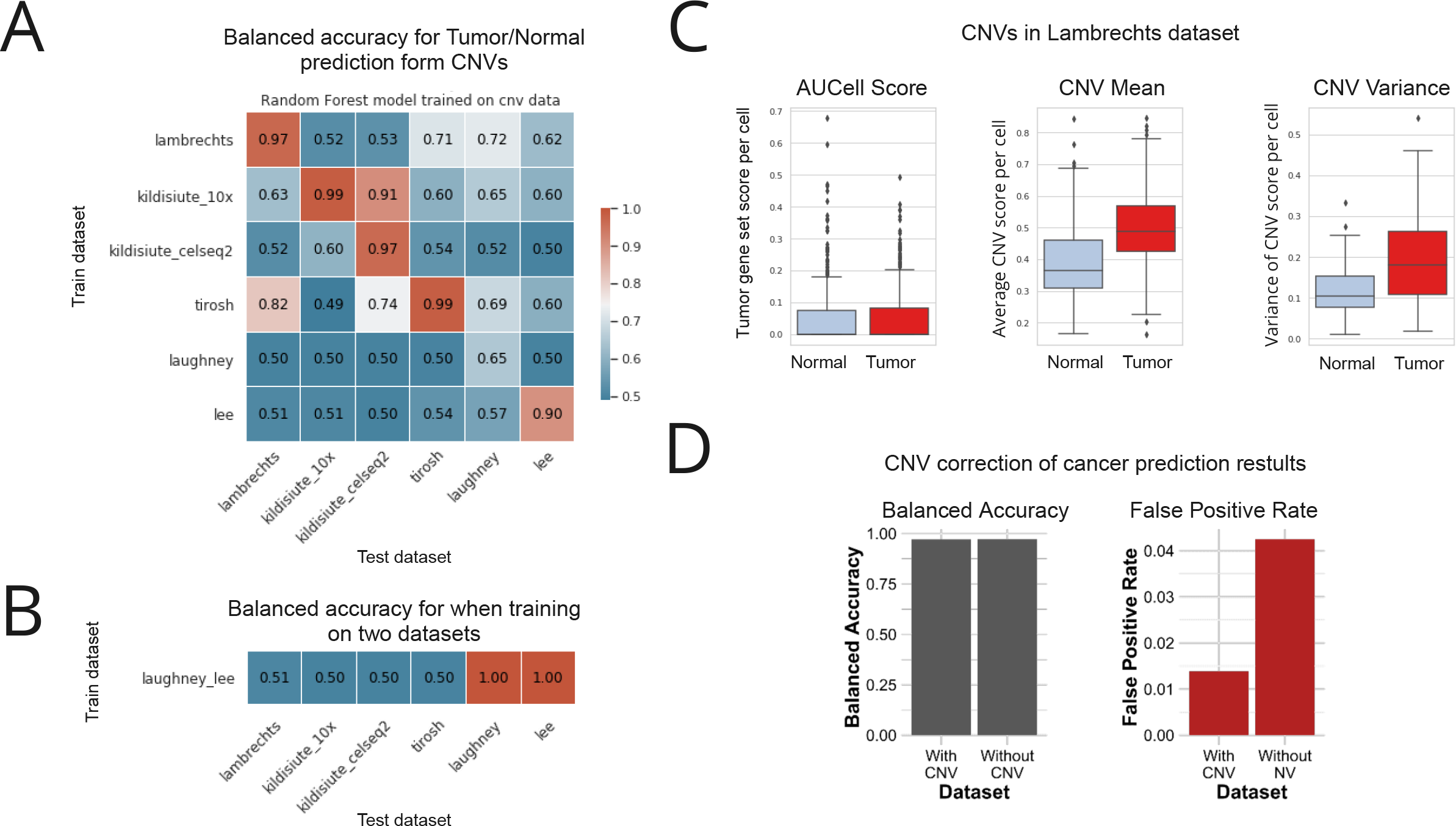
Multio-mics decreases the false positive rate. A) Balanced accuracy of random forest classifiers trained on each of the tumor types. The classifiers have excellent performance on the same samples they were trained on, or on similar tumors (such as the Kildisute et al. neuroblastoma sample), while they fail to generalize to other tumor types. B) Training on multiple tumor types does not improve the generalization of the classifiers. C) Ikarus misclassified cells can be discriminated based on the average CNV and variance of CNV values. Tumor cells misclassified as normal cells have significantly higher values of both the average CNV and the variance of CNV, than the corresponding normal cells. D) Integration of the CNV proofreading decreases the false positive rate from 4% to 1%, with the same average balanced accuracy.

## Discussion

We have implemented a two step approach for solving a problem that is perceived as simple:discriminating tumor cells from normal cells. In the first step, Ikarus pipeline integrates multiple expert labeled datasets to extract gene sets which discriminate tumor cells from normal cells. In the second step, Ikarus uses a robust gene set scoring along with adaptive network propagation for cell classification. By using robust gene set scoring and network propagation we have mitigated two common problems in single cell analysis: the influence of batch effects on sample comparison and parameter optimization during clustering.

The effect of technical differences between single cell datasets is usually resolved using integration methods. Single cell integration methods require extensive tuning of sets of parameters, most of which have non-intuitive effects on the results. Moreover, the accuracy of the resulting integrations can not be trivially evaluated without extensive usage of biological priors. Gene set scoring methods are robust, because they use “within sample” rank based scores instead of direct comparison of measured expression values between different samples. The only technical variable that influences gene set scoring is the percentage of genes from the gene list which are detected in each cell. We have however extensively tested the influence of the number of genes on the classification accuracy.

A common step in single cell analysis is aggregation of cells into clusters, which are then used for cell type annotation. Clustering is, however, a procedure with an inherently high number of parametric options. It is extremely hard, if not impossible, to choose a set of parameters that would produce the same level of accuracy (same cell types) on different datasets, which often necessitates manual intervention to deduce the best clustering resolution. Because cell types and cell states form highly connected modules within the cell - cell similarity graph, we have therefore opted to replace clustering with network propagation. Network propagation is a procedure where the uncertainty of cell annotation can be reduced by integrating the annotation score of each individual cell with the scores of its nearest neighbours. Network propagation represents a parameterless alternative to clustering, while retaining the same level of sensitivity for cell annotation.

By exploring a multi-omics approach, we have tried to increase the accuracy of the normal - tumor cell discrimination. Using inferred CNVs, we have shown that the information from copy number variations does not generalize across different cancer types. By including the copy number variation as a proof reading step, we reduced the false positive rate of the classifiers. It is still an open question, though, by how much would single cell multi-omic measurements improve the classification (for example, by concurrently measuring mutations, CNVs, chromatin accessibility, and expression in the same single cells). Currently, such methods are either in their infancy, and the required data is not available, or have a very limited profiling range (profile only a handful of loci).

Ikarus is currently constrained by the reliance on well annotated single cell datasets. For both the gene set definition and testing, we rely on expert provided cell annotation. This requirement has limited our training and testing capabilities to the handful of profiled, and annotated cancer types. However, the exponentially increasing number of single cell data sets will enable us to increase both the number of training datasets, as well as to test Ikarus on currently unavailable tumor types, for instance, soft tissue sarcomas. Moreover, the increasing quality of single cell data sets, most importantly, increasing gene coverage, will also increase the utility of Ikarus as a gene set based classification method.

By integrating multiple datasets we have derived a tumor signature gene list which is surprisingly refractory to annotation. The gene list contains a sub-module (n = 34), that encompasses genes involved in the cell cycle. All of the other genes, however, showed little modularity, and a lack of enrichment in any single annotation category. Interestingly, the genes were highly expressed in all of the available PDX and CCLE cancer models. The ablation studies showed that the classifier was robust to the removal of any one of the genes. The low co-expression, combined with the lack of sensitivity to the gene removal indicates that the tumor signature genes provide mutually synergistic information towards the classification accuracy. Ikarus classifier, however, is not limited to tumor cell detection. It can be used to detect any cellular state, such as cell types. The only requirements are that the cellular state is present in at least two independent experiments, which are expertly annotated.

Automatic, parameterless discrimination between tumor cells and the surrounding tumor-associated tissue has multifactorial utility. Tumor cells can be streamlined into algorithms for neoepitope prediction, thereby enabling direct, clinically relevant insights. Furthermore, the increasing availability of multi-omics measurements would enable automatic genetic characterization of tumor subpopulations, and the subpopulation based recommendation of best therapeutic course of action. Application of automatic tumor classification on spatial sequencing datasets enables direct annotation of histological samples, thereby facilitating automated digital pathology.

The current scale of development in single cell biology (on both the technological and computational levels) shows promise for quantitative characterization of the complete tumor heterogeneity, for each individual. However, before the personalized medicine approach can be readily adopted, every step in the data analysis needs to be completely automated, with robust performance guarantees. Ikarus pipeline represents one step towards the implementation of personalized cancer therapy.

## Datasets

### Single cell RNA-seq data

Gene expression values for single cell RNA-seq experiments are available through the corresponding publications. If not explicitly declared otherwise, the 10X genomics protocol was used for scRNA-seq. Laughney et al. provide a lung adenocarcinoma (primary tumors and metastases) data set that include 40505 cells coming from 17 patients. For our purpose 39414 cells are considered tumorous and 1091 are normal. This dataset serves as input for model building. 63689 cells from 23 colorectal cancer patients are coming from Lee et al. 16248 cells are considered tumorous and 47441 are normal. These cells are used as input for model building. A Non-small-cell lung cancer data set is coming from Lambrechts et al. It considers 52698 cells, of which 7447 are tumorous and 45251 are normal. This data set is used for model testing. Puram et al. published 5578 single cells from 18 head and neck squamous cell carcinoma patients. They performed Fluorescence-activated cell sorting for scRNA-seq. 2215 cells are tumorous, 3363 cells are normal. We use this data set for model testing. Kildisiute et al. published a neuroblastoma cell atlas. We used 6442 cells (10X) from 5 patients (1766 tumorous, 4676 normal) and 13281 cells (CEL-seq2) from 16 patients (1630 tumorous, 11651 normal) as two distinct data sets for model testing (Table S1 - Sheet 2).

### ENCODE cell line dataset

Gene expression values for primary cells, cell lines, and cancer cell lines were downloaded in batch from the ENCODE portal with the following query: Assay title: “polyA plus RNA-seq”; Status: “released”; Perturbation: “not perturbed”; Organism: “Homo sapiens”; Biosample classification: “cell line”, “primary cell”; Genome assembly: “GRCh38”. Identifiers corresponding to the acquired data totalling 860 files are provided in the supplementary materials (Table S1 - Sheet 4). Downloaded expression tables were merged and standardized by a custom R script prepare.data_encode.R to a combined gene expression matrix that includes all input data, HGNC symbol gene annotation and cell annotations. For gene expression quantification log2(TPM) with a pseudocount of 1 was used. Based on those components an AnnData object is created which is then provided as an input to Ikarus package.

### Microdissection dataset

The gastric cancer microdissection dataset comprises laser capture microdissected (LCM) stromal and cancer regions collected from a patient cohort (n = 8) totalling 16 samples. Microdissected tissue for each sample was pooled together before library preparation to account for the absence of replicates. Gene expression quantification of stromal and cancer samples, as provided by the authors of the study [28] in the form of raw counts, was first standardized to Ikarus format and then used as an input to Ikarus pipeline.

### Databases

- Human protein atlas (https://www.proteinatlas.org/humanproteome/pathology) [42]
- Prognostic genes [43]
- Gene fusion (ChiTaRs) [44]
- SEEK (co-expression database) (https://seek.princeton.edu/seek/) [39]
- g:Profiler (https://biit.cs.ut.ee/gprofiler/) [41]
- CancerSEA (http://biocc.hrbmu.edu.cn/CancerSEA/home.jsp) [38]
- MsigDB (GO, Hallmark gene sets) [40]
- Atlas of co-essential modules [47]
- DepMap Achilles scores (https://depmap.org/portal/download/) [48]
- COSMIC (cancer.sanger.ac.uk) [49]

### Code Availability

Ikarus is a python package that can be found on the following link: https://github.com/BIMSBbioinfo/ikarus

Code for reproducing the figures can be found on the following link: https://github.com/BIMSBbioinfo/ikarus---auxiliary

## Methods

### Ikarus workflow

The presented Ikarus pipeline consists of three modules. First step is the gene selection and integration that detects cell group specific gene sets. Next, based on the constructed gene sets, unknown cells of interest are scored and annotated. While the first step is optional, as users can provide their own gene lists, the latter steps are mandatory to make a prediction. For making predictions Ikarus’ handling is opted to be close to the scikit-learn workflow, which means 1. load data, 2. initialize a model, 3. fit the model and 4. make the actual predictions on unknown data. In general, annotated data objects are used as data format (AnnData, https://anndata.readthedocs.io). Each individual step is described in more detail in corresponding subsections below.

### Defining gene signature lists

The count matrices of the input AnnData objects are normalized to the total number of reads per cell and log transformed with a base of 2 and a pseudocount of 1. In addition to that, each AnnData object must contain for each cell the corresponding cell type in the observation section, possibly in multiple columns for multiple hierarchical cell type levels (e.g. tumor and normal cells, or tumor, epithelial and immune cells). Then, for each gene in the input data set a t-test with overestimated variance is used to compute an approximation of log 2 fold change between two cell type classes, one up-regulated and one down-regulated class. Those classes are provided by the user and should be chosen in accordance with the considered columns of the AnnData observation section. Users can either perform only one comparison (e.g. tumor vs. normal cells) but also multiple (e.g. tumor vs. epithelial cells and tumor vs. immune cells). This is done independently for each dataset. For each gene and for each pair-wise comparison, the associated log 2 fold changes (if p_adj_ < 0.1, neglected otherwise) of the different input data sets are averaged. According to these average values the genes are then sorted, highest to lowest and a user defined number of top genes is selected. The final list of upregulated genes is derived by taking either the intersection or the union of selected genes across all of the comparisons. The whole procedure is performed once for the case that the class of interest (e.g. tumor) is up-regulated (here we take the intersection of selected genes across all of the comparisons) and once for the case that this class is down-regulated (here we take the union of selected genes across all of the comparisons). That way, we obtain two final gene sets. One set representing genes enriched in the class of interest (i.e. enriched in tumor cells), and a set depleted in the class of interest (depleted in tumor cells).

### Cell scoring using gene sets

Both the tumor and normal gene sets were used to score each of the cells in each of the experiments using AUCell [50]. As input to AUCell we provide the gene expression matrices that were normalized to the total number of sequenced reads per cell, and subsequently transformed using the log2(x + 1) function. AUCell requires that the dataset contains at least 80% of the genes from the input gene set.

We have noticed that the AUCell scores do not behave properly in some of the bulk sequencing data sets. Namely, samples which had similar transcriptomes sometimes had widely different AUCell scores. We have therefore implemented an option to use a different gene set scoring function - ssGSEA.

### Logistic classifier training

A logistic classifier was trained on the combined Lee et al. and Laugney et al. datasets. Scores of normal and tumor gene signatures were used as the input, and the tumor/normal class assignment as the target variables.

### Cell annotation using network propagation

Ikarus implements the cell annotation as an iterative two step process of cell type assignment and label propagation. In each iteration we assign labels to cells with a decreasing stringency threshold, which are then propagated to their nearest neighbours. Firstly, the labels are assigned to the most probable cells, based on a robust stringency threshold.ells below the stringency threshold have their LR probabilities masked to zero. The stringency threshold is defined based on the order statistic of the gene set score difference between the two classes of interest. In the first iteration it is the 90% percentile of the (tumor - normal) gene set score difference. The label propagation is then obtained by computing the dot product of neighborhood connectivities and LR class probability estimates. Annotations are derived from the propagated class probabilities. Within each iteration step, the stringency threshold is reduced using exponential decay:

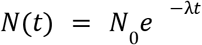

Where

*N*_0_ is the starting stringency threshold;

*t* is an iteration step;

λ is an exponential decay constant.

The cell - cell graph, used for label propagation, is constructed using the normal and tumor gene signatures according to [51], as adopted in [52].

The label propagation procedure is repeated until less than 0.1% of cell annotations change.

## CNV correction

### Classification improvement using copy number variations

To improve the classification results we used inferred CNV scores as an additional source of information. inferCNV [46] was used to compute CNV scores. A cut-off value of 0.1 was chosen for gene selection. CNV prediction was performed via HMM. For tumor sub-clustering the parameters were kept default (hclust = ‘ward.D2’, tumor_subcluster_pval = 0.05), though tumor_subcluster_partition_method was set to ‘qnorm’, as this is claimed to be reasonable faster than ‘random_trees’. No prior information on distinct clusters was provided. In a self-supervised fashion, we used the current Ikarus cell annotations as pseudo-labels to train an additional logistic regression model (LR). The LR takes as its input per cell inferred CNV values, and predicts the cell annotations. The LR itself is trained on all cells from the validation dataset, e.g. Lambrechts et al., Kildisiute et al., Puram et al. This model was then used to make predictions on the same dataset assuming that Logistic Regression should not overfit on this task. The outcome is then considered as the final corrected Ikarus prediction.

## Gene set characterisation

### Comparison with published gene sets

The tumor signature gene set assembled in this study was characterized by comparison with publicly available gene sets provided by multiple public resources. We considered gene sets of various provenances, e.g. all Homo sapiens gene sets published by MsigDB [40], cancer specific gene sets that represent distinct functional tumor states (CancerSEA) [38], novel gene lists covering previously unidentified members of cellular signalling pathways [47].

Gene sets as provided by MsigDB (n = 31120) were assessed via the interface provided by the R package msigdbr version 7.2.1. Next, Intersections and unions of the Cancer gene set (this study) with every human gene set from that release (version 7.4) of MsigDB, as well as the members of the intersections and original sizes of query gene sets, were computed. CancerSEA resource provides a collection of functional cancer gene sets derived from a multitude of single cell studies, thereby supplying a single cell level scope to the cancer functional hallmarks. For characterization of the Cancer gene set assembled in this study, we downloaded 14 gene sets n_min_ = 66, n_max_ = 201 from the CancerSEA resource (http://biocc.hrbmu.edu.cn/CancerSEA/goDownload) representative of distinct cancer functional states and intersected it with our gene set. The results of this analysis were presented with the UpSet plot framework [53] implemented in the ComplexHeatmap R package [54], mode intersect.

To account for recent advances in the annotation of cellular signaling pathways, we downloaded a novel collection of gene sets composed of genes previously unmapped to any signal transduction pathway. Namely, we acquired 11 gene sets of various sizes n_min_ = 10, n_max_ = 164) and intersected with the tumor signature gene set of this study. The visualization of this analysis was similar, i.e. the intersections were presented with an UpSet plot implemented in ComplexHeatmap R package, mode intersect.

### Gene fusions

Data on human gene fusions were downloaded from the ChiTaRS resource as was provided on 2019/08/16 (http://chitars.md.biu.ac.il/index.html) [44]. First, we constructed a background distribution from randomly selected sets of genes that were expression-matched to the tumor signature gene set (this study). Every random gene set was intersected with fused genes from the database and the resulting intersection sizes were used to fill a background distribution. Lastly, the tumor signature gene set from this study was intersected with the list of fused genes to compare with the background distribution. This analysis was done separately on 5’- and 3’-fused genes.

### Co-expression analysis

To investigate genes that are co-expressed with the tumor signature gene list across many datasets, we took advantage of web-based platform SEEK (https://seek.princeton.edu/seek/) [39]. We queried the Cancer gene list to the SEEK search engine and downloaded a SEEK-generated ranked list of co-expressed genes. For further analyses we used top 150 genes from the ranked list.

### Gene Ontology (GO) analysis

The GO analyses throughout this study were done using the framework provided by gprofiler2 R package [41]. From the default run settings, p-values threshold was changed to 10e-4 and correction_method option set to g_SCS.

### Gene set sensitivity testing

To measure Ikarus’ robustness on the extracted tumor gene list, we performed the following analyses:

### Gene set size

Using the Lambrechts et al. lung cancer dataset, we iteratively computed Ikarus’ balanced accuracies ablating a random section of the tumor gene list in a cumulative step-wise manner before scoring and prediction steps. Namely, we randomly removed 20, 40, 60 and 80% percent of the tumor gene list. Every ablation percentage was itself reiterated 20 times. The predictions were not CNV-corrected.

### Single gene ablation

Further, we investigated the prediction value of individual genes in the gene list. We employed a similar procedure as before, but in contrast to ablating the whole sections of the list, we removed an individual gene from the list per iteration before computing Ikarus’ balanced accuracies.

### CNV Analysis

To investigate the overrepresentation of copy number amplifications among the genes in the extracted tumor gene list we referred to the COSMIC database. Namely, we downloaded a complete COSMIC collection of copy number alterations and stratified it by tumor tissue of origin (n = 27). Next, we iteratively intersected the tumor gene list from this study with significantly amplified genes (denoted as “gain” in the COSMIC table) over tumor tissues of origin. As a random control, we prepared similarly sized random gene sets (n = 162) that were expression-matched to the original tumor gene list. Expression matching was done on Laughney et al., Lee et al., and Lambrechts et al. datasets independently. In total, 150 random gene sets were generated, 50 sets per expression dataset.

## Figures

**Figure S1.**
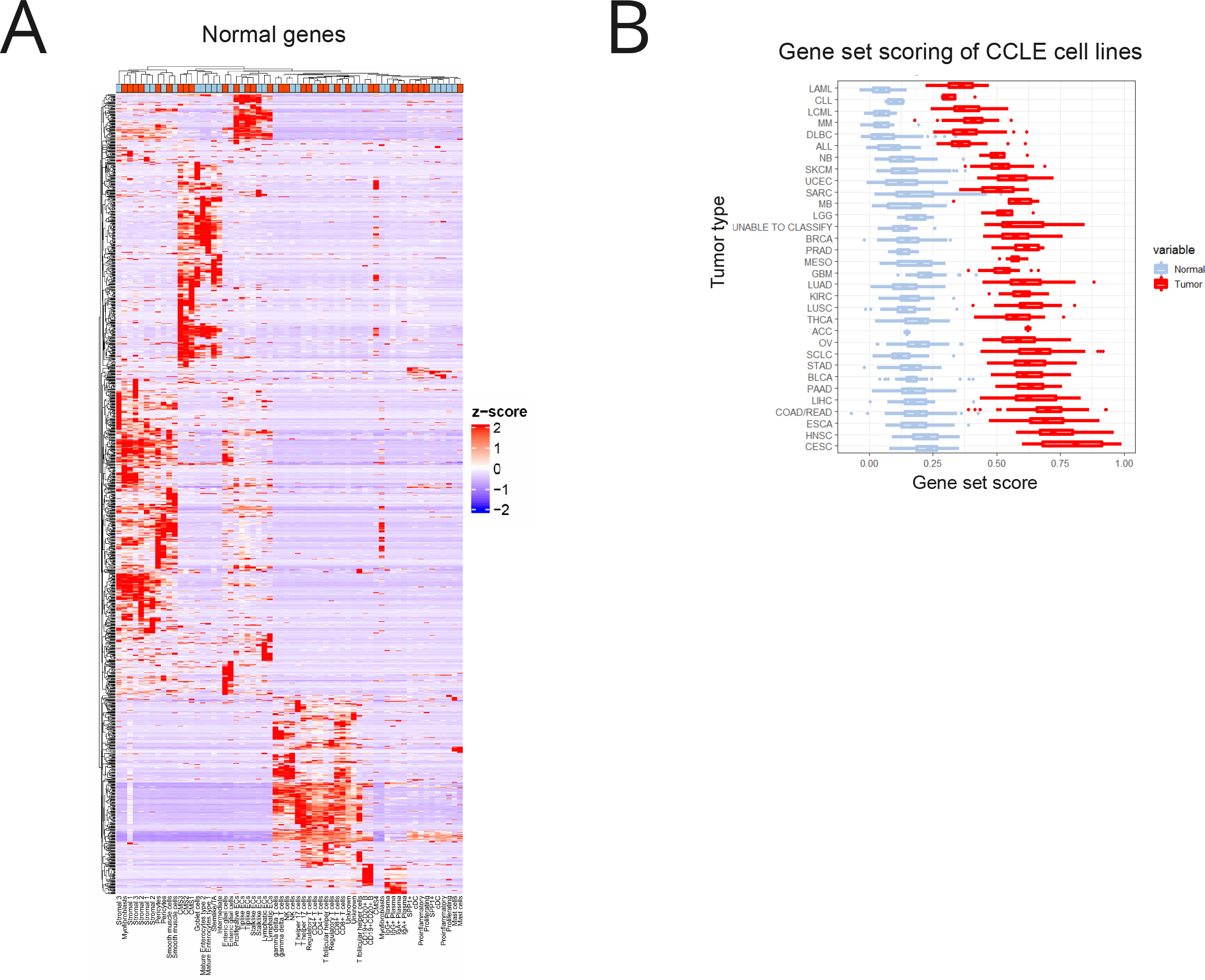
Tumor and normal gene signature characterization. A) The normal gene signature list visualized on the Tirosh dataset (which was not used in the gene list definition). Normal gene signature captures mostly cell type specific gene expression. B) Tumor and normal gene signature scores of the cancer cell line encyclopedia (CCLE) data. Tumor gene signature shows a significantly higher score distribution in all cancer types present in the dataset.

**Figure S2.**
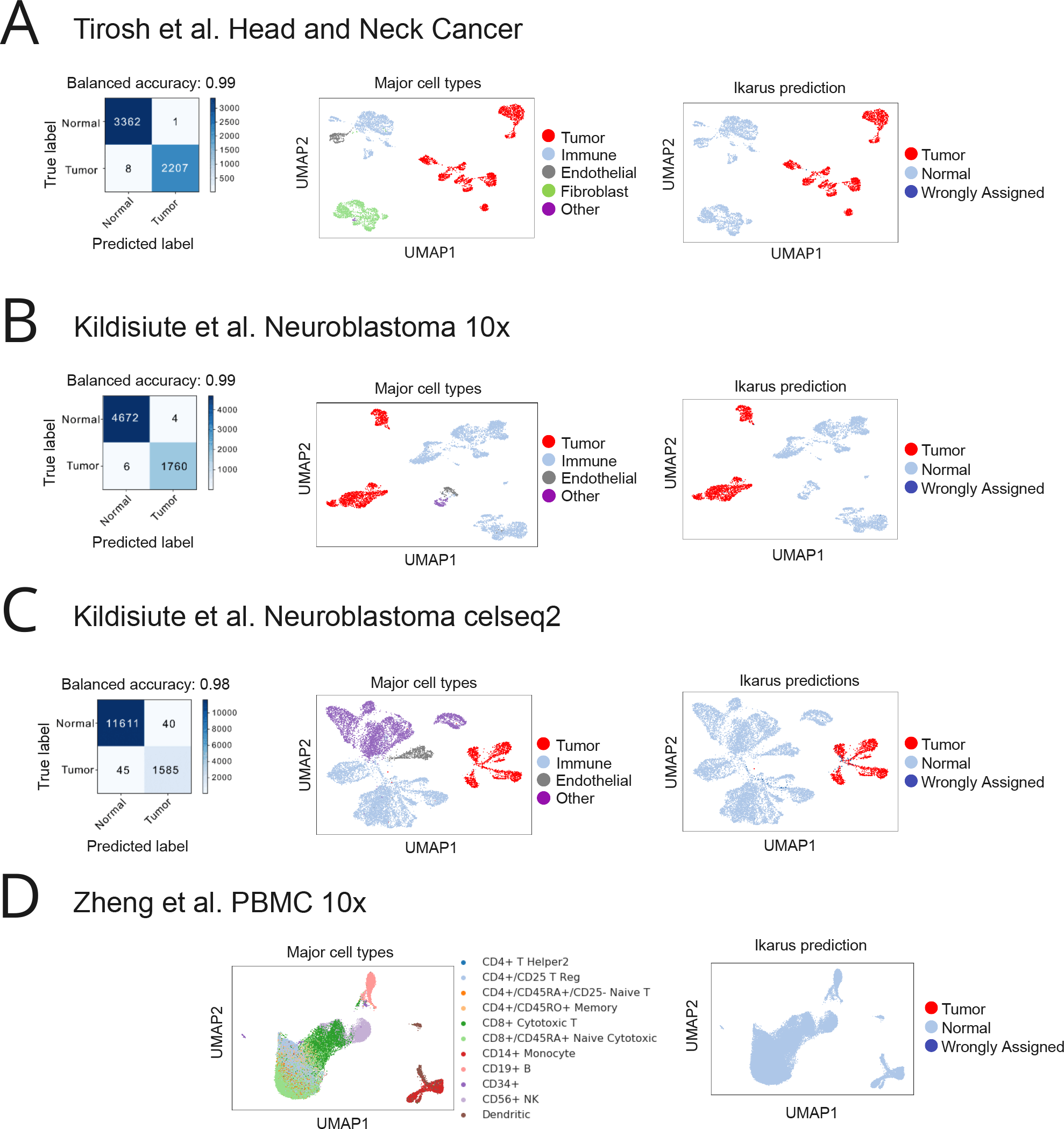
Ikarus performance on multiple test data sets. A) Performance of Ikarus classifier on the Puram et al. Head and Neck Cancer dataset B) Performance of Ikarus classifier on the Kildisiute et al. Neuroblastoma dataset sequenced with 10x. C) Performance of Ikarus classifier on the Kildisiute et al. Neuroblastoma sequenced with CEL-Seq2. D) Ikarus correctly recognizes all cells from a healthy peripheral blood as non-tumorous.

**Figure S3.**
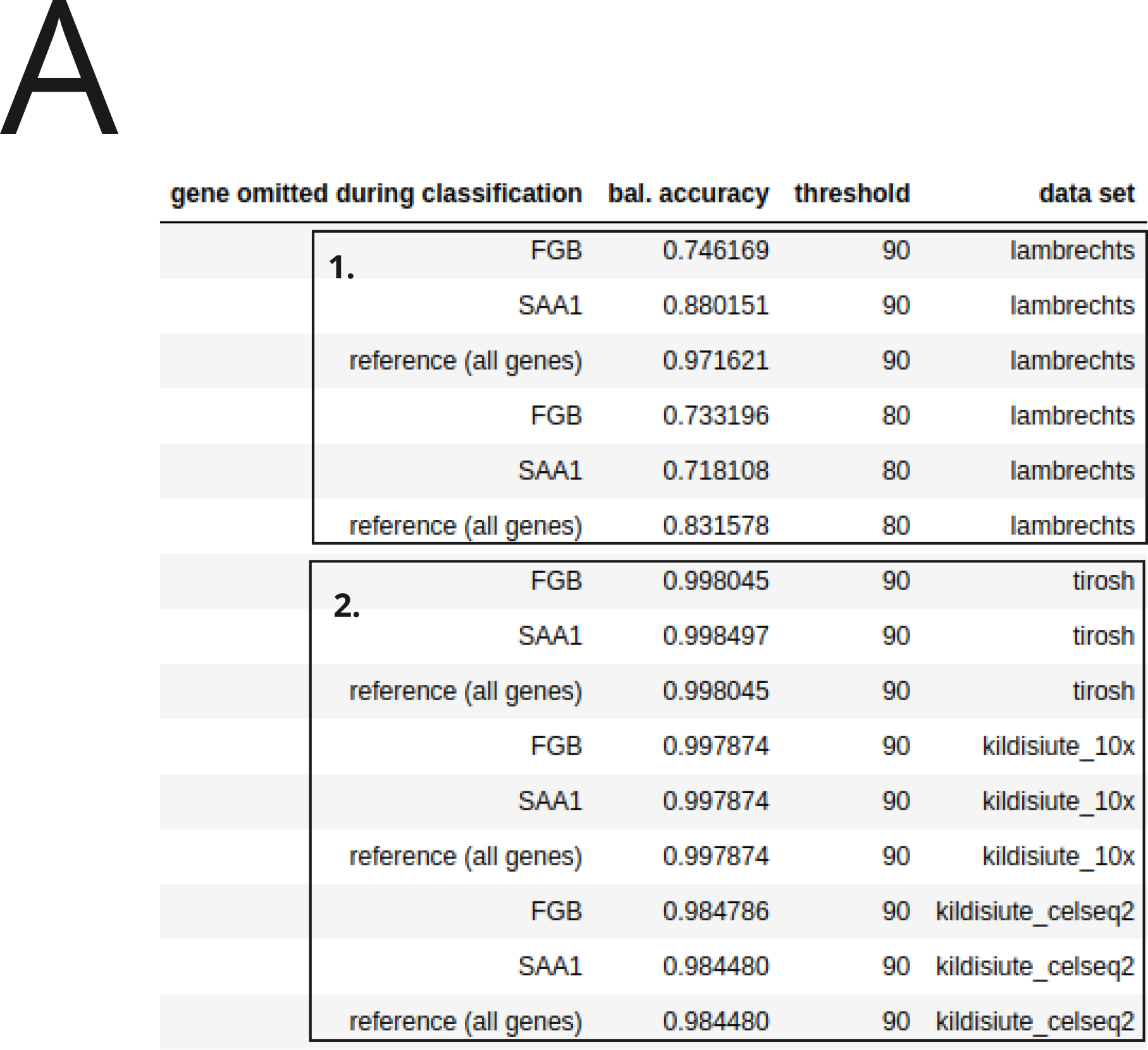
Effects of SAA1 and FGB ablation on predictions. A) Balanced accuracy of Ikarus’ predictions after one by one ablation of FGB and SAA1 in lung, head and neck, and neurological (neuroblastoma) cancers.

**Figure S4.**
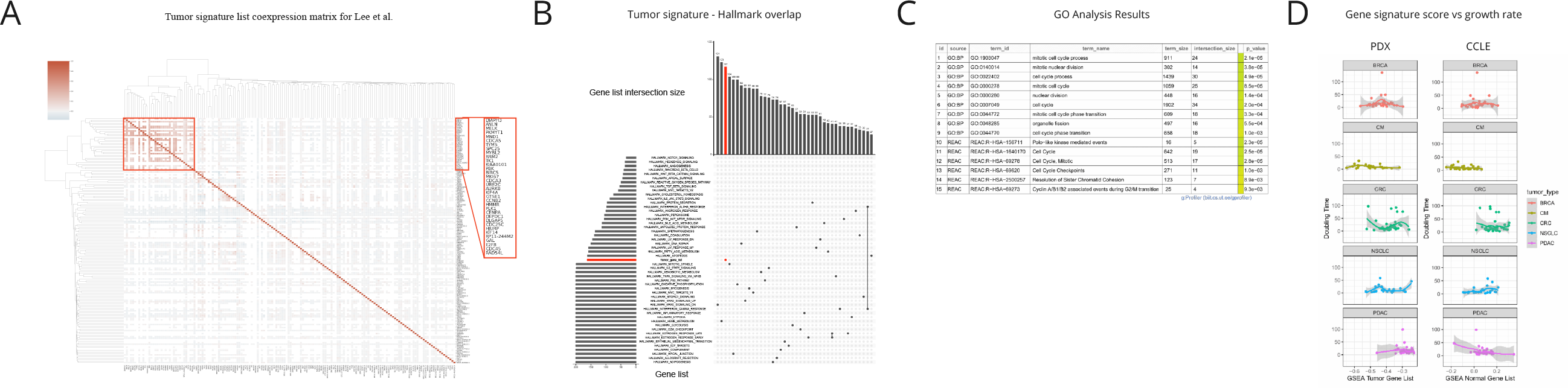
Additional annotation of the tumor gene signature. A) Correlation analysis of the tumor gene signature from Lee et al. Genes belonging to the cell cycle module are marked with a red rectangle. B) Tumor gene signature shows limited overlap with most of the hallmark gene sets. Only intersections of size 27 and more are shown. C) Gene set overrepresentation analysis using gProfiler. D) Relationship between the tumor gene signature scores and the growth rate of PDX and CCLE models. There is limited correlation between the growth rate and the gene signatures.

## Acknowledgments

We would like to sincerely thank Florian Uhlitz for a very constructive discussion and critical reading of the manuscript.

## Author contributions

AA conceptualized and planned the project. JD and AB jointly executed all of the computational analyses. BU, and VF wrote the manuscript, with comments from JD and AB. JR critically edited the manuscript. BU, VF, and AA jointly supervised the work. All authors read, edited, and confirmed the final manuscript. AA acquired funding and supervised the project.

